# Structural neuroimaging signatures of anorexia nervosa features in a mixed sample enriched for disease vulnerability

**DOI:** 10.1101/498097

**Authors:** Amy E. Miles, Allan S. Kaplan, Yuliya S. Nikolova, Aristotle N. Voineskos

## Abstract

Brain-behavior relationships that could provide insight into risk-associated pathophysiology have not been thoroughly assessed in anorexia nervosa (AN). Therefore, we sought to identify grey and white matter signatures of AN symptoms and risk factors (trait anxiety, set-shifting impairment) in a sample enriched for AN vulnerability, including acute and remitted AN patients and their unaffected sisters (*n* = 72, aged 18 – 48 years). MRI/DTI data were acquired on a 3T scanner and processed with Freesurfer and FSL TBSS. Relationships between clinical variables of interest and regional subcortical volume, vertex-wise cortical surface architecture (thickness, surface area, local gyrification), and voxel-wise white matter microstructure (FA, MD) were tested with separate linear regressions, including age, BMI, lifetime AN diagnosis, and intracranial volume as covariates, where appropriate. Significance was determined using a Bonferroni-corrected threshold, *p*(*t*) ≤ 0.001. We detected distinct associations linking AN symptoms to lateral occipital cortical thickness and insular/cingulate gyrification and trait anxiety to lingual cortical thickness and superior parietal gyrification, and we detected overlapping associations linking AN symptoms and set-shifting impairment to frontoparietal gyrification. No other brain-behavior relationships emerged. Our findings suggest that variations in site-specific cortical morphology could give rise to core features of AN and shared temperamental and cognitive-behavioral risk factors for AN.

## INTRODUCTION

Anorexia nervosa (AN) is a complex psychiatric disorder marked by persistent restriction of energy leading to severely low body weight and characterized by strong drive for thinness and overwhelming body dissatisfaction (for a review, see Treasure et al., 2015). AN typically manifests in adolescent females, and it is highly heritable (Bulik et al., 2006). Without evidence-based treatments for adult AN, mortality is high, and fewer than half of patients achieve full recovery.

Clinical neuroimaging studies have identified neuroanatomical differences between patients with AN and controls that may provide partial insight into disorder pathophysiology (for a review, see Seitz, Herpertz-Dahlmann & Konrad, 2016). Specifically, prior T1-weighted magnetic resonance imaging (MRI) research suggests case-control differences in subcortical volume, cortical thickness, cortical surface area, and local gyrification, while diffusion tensor imaging (DTI) studies suggest variations in white matter microstructure. Given strong evidence of additive genetic effects and consistent identification of neurodevelopmental risk factors for AN (e.g. perinatal complications, childhood trauma), it is possible that a substantial proportion of the observed case-control differences in brain structure may be part of a vulnerability profile that predates disorder development. However, the interpretation of these results is complicated by starvation-driven pathophysiology (e.g. fluid and electrolyte imbalance, endocrine dysregulation), effects of which cannot be readily distinguished from premorbid differences in brain structure (Treasure et al., 2015).

Identifying differences in brain structure between acute and remitted AN patients can help disentangle state-dependent neuroanatomical effects from their trait-like counterparts. Several such studies from our group and others have revealed that the extent to which the aforementioned variations in brain structure are driven by acute malnutrition appears phenotype-dependent; whereas reductions in subcortical volume and cortical thickness appear to normalize with weight-recovery, indicating state-dependence, in some cases, changes in local gyrification and white matter microstructure appear to persist into long-term remission, suggesting trait-relevance (Miles, Voineskos, French & Kaplan, 2018; Miles, Kaplan, French & Voineskos, 2019; Seitz et al., 2016).

Beyond case-control comparisons, mapping structural brain changes onto specific dimensions of AN-related psychopathology can illuminate the biological basis of distinct, tractable facets of the otherwise complex AN clinical manifestation. Within this framework, testing relationships between neuroanatomical phenotypes and measures of psychopathology that are unique to AN and shared across psychiatric disorders is critical since AN rarely manifests in isolation; up to half of AN patients report one or more comorbid anxiety-spectrum disorders (Kaye, Bulik, Thornton, Barbarich & Masters, 2004), and significant genetic correlations between AN and obsessive-compulsive disorder (OCD) have been reported (Anttila et al., 2016). Given that shared genetic risk could contribute to temperamental and cognitive-behavioral hallmarks of AN that also manifest in anxiety and OCD (for a review, see Kaye, Wierenga, Bailer, Simmons & Bischoff-Grethe, 2013), including measures of trait anxiety and cognitive flexibility alongside measures of core AN symptomatology can provide important complementary insight into AN pathophysiology.

While prior case-control studies have compared acute and remitted AN patients in order to disentangle state- and trait-like neuroanatomical abnormalities, dimensional mapping across the full spectrum of psychopathology in a sample enriched for AN vulnerability has not been undertaken. To fill this important gap, here we aim to comprehensively assess brain-based correlates of disease risk by testing associations between core AN symptomatology and shared temperamental and cognitive-behavioral risk factors for AN (i.e., trait anxiety and set-shifting impairment), dimensional measures that can vary widely across individuals and disease states (Pollice, Kaye, Greeno & Weltzin, 1997; Roberts, Tchanturia & Treasure, 2010), and distinct neuroanatomical phenotypes that have been implicated in disease vulnerability and manifestation (i.e., subcortical volume, cortical thickness, cortical surface area, local gyrification, and white matter microstructure). In order to capture brain-behavior relationships across the full spectrum of disease risk, we will test these associations in a mixed sample, including women with low suspected vulnerability to AN (i.e., unrelated healthy controls), women with high suspected vulnerability to AN but no disease manifestation (i.e., unaffected sisters), women with a history of disease manifestation (i.e., remitted patients), and women with current disease manifestation (i.e., acute patients).

Although this study is exploratory, we have several preliminary hypotheses. Above all, across groups we expect to observe negative dimensional associations between measures of core AN symptomatology, trait anxiety, and set-shifting impairment and neuroanatomical phenotypes that appear compromised in prior case-control studies of AN. The latter includes subcortical volume, cortical thickness, cortical surface area, local gyrification, and fractional anisotropy (FA), an index of white matter microstructure and putative measure of fiber integrity (Alexander, Lee, Lazar & Field, 2007; Leppanen, Sedgewick, Cardi, Treasure & Tchanturia, 2019; Miles et al., 2018; Seitz et al., 2016). We also expect to observe positive associations between the aforementioned clinical variables and mean diffusivity (MD), an index of white matter microstructure and putative measure of cellularity (Alexander et al., 2007), increases in which have been reported in AN (Miles et al., 2019).

## METHODS AND MATERIALS

### Sample recruitment

Data was collected from adult females (*n* = 72, aged 18 – 48 years) recruited from university campuses and hospital and community-based clinics in the Greater Toronto Area. The sample included women with acute AN (acAN, *n* = 24), women with remitted AN (recAN, *n* = 24), unaffected sisters of acAN/recAN participants (sibAN, *n* = 12), and unrelated healthy controls (HC, *n* = 12). Participants were medically fit for imaging, they did not meet criteria for a substance use disorder, and they reported no lifetime psychosis, head trauma, or neurological problems.

Group-specific inclusion criteria were as follows: (1) acAN: current AN diagnosis, BMI < 18.0 kg/m^2^; (2) recAN: past AN diagnosis, normal weight for 12+ consecutive months, symptom absence for 12+ consecutive months; (3) sibAN: normal weight, no lifetime ED diagnosis; (4) HC: normal weight, no lifetime psychiatric history. For all groups, normal weight was defined as BMI > 18.5 kg/m^2^ and BMI < 30.0 kg/m^2^.

The study was approved by the Centre for Addiction and Mental Health Research Ethics Board. All participants gave written informed consent.

### Clinical assessment

Participants were assessed with the following measures. This battery was designed to index deficits frequently associated with AN.

1. a semi-structured interview, administered by the first author, was used to collect demographic and clinical information;
2. the Structured Clinical Interview for Diagnosis, Research Version (First, Spitzer, Gibbon & Williams, 2002), modified to reflect guidelines from the Diagnostic and Statistical Manual of Mental Disorders, Fifth Edition (American Psychiatric Association, 2013), was used to confirm AN diagnosis and screen for psychosis and substance use;
3. the Eating Disorder Examination Questionnaire, Version 6.0 (Fairburn & Beglin, 2008) was used to confirm AN diagnosis;
4. the Eating Disorder Inventory-3 (Garner, 2004) was used to assess core AN symptomatology, indexed using drive for thinness, bulimia, and body dissatisfaction scores;
5. the State-Trait Anxiety Inventory (Spielberger & Gorsuch, 1983) was used to assess trait anxiety; and
6. the WCST-64: Computer Version 2-Research Edition (Kongs, Thompson, Iverson & Heaton, 2000) was used to assess set-shifting impairment, indexed using number of perseverative responses.

### Image acquisition and processing

MRI data was collected on a 3T scanner (Echospeed; General Electric Medical Systems) at the Centre for Addiction and Mental Health. High-resolution, 3D T1-weighted structural scans were acquired using a fast-spoiled gradient-recalled echo sequence with the following parameters: TE = 3ms; TR = 8.2ms; TI = 650ms; *α* = 8°; FOV = 24cm; NEX = 1; 0.9mm isotropic voxels, no gap. For DTI, sixty diffusion-weighted (*b* = 1000 s/mm^2^) and five baseline scans (*b* = 0 s/mm^2^) were acquired using an echo planar imaging sequence with dual spin-echo and the following parameters: TE = 88 ms; TR = 8800ms; FOV = 256mm; 128×128 encoding steps; 2.0mm isotropic voxels, no gap.

Analysis of MRI/DTI data was carried out with tools from the Freesurfer Software Suite (http://surfer.nmr.mgh.harvard.edu, version 6.0.0) and FMRIB Software Library (https://fsl.fmrib.ox.ac.uk, version 5.0.10). Processing pipelines are described in detail in Miles et al. (2018, 2019). Local gyrification was quantified using the local gyrification index (LGI) (Schaer et al., 2012).

### Regression analyses

Main effects of core AN symptomatology, trait anxiety, and set-shifting impairment on MRI/DTI-derived measures were tested with separate linear regressions including age, BMI, and lifetime AN diagnosis as nuisance variables. Total intracranial volume was also included as a nuisance variable in analyses of subcortical volume and cortical surface area. Each regressor was standardized prior to analysis, and age and clinical variables of interest were square root-transformed to reduce skewness. All primary analyses were performed sample-wide.

When regional subcortical volume was the dependent variable of interest, significance was determined using a Bonferroni-adjusted threshold, *p*(*t*) ≤ 0.001, and analyses were repeated for each of the following segmentations: accumbens area, amygdala, caudate nucleus, hippocampus, pallidum, putamen, and thalamus. To reduce the number of comparisons, volumes were averaged over left and right hemispheres.

When vertex-wise cortical thickness, cortical surface area, or local gyrification index was the dependent variable of interest, significance was determined using Monte Carlo simulation with 10,000 repetitions and vertex-wise and cluster-forming thresholds, *p*(*t*)_FWER_ < 0.05. When voxel-wise FA or MD was the dependent variable of interest, significance was determined using permutation testing with 10,000 repetitions and threshold-free cluster enhancement. For vertex/voxel-wise analyses, clusters were retained if *p*(*t*)_FWER_ ≤ 0.001.

### Post hoc analyses

In order to ascertain if observed main effects of clinical score on MRI/DTI-derived measures were driven by particularly strong relationships in one or more participant subgroups (acAN, recAN, sibAN, HC), we tested within-group associations between relevant clinical scores and region or cluster means using Pearson correlation.

In order to further probe robustness of observed sample-wide correlations, we conducted k-fold cross validation and leave-one-group-out cross validation. These resampling procedures allow the user to evaluate model fit by sequentially excluding random groups of participants or designated groups of participants, respectively. For the latter, we computed prediction errors based on sequential testing of linear models excluding one participant subgroup at a time. All post hoc analyses were run in RStudio (RStudio Team, version 0.99.489).

## RESULTS

### Sample characteristics

Sample characteristics are summarized in Tables 1 and 2 and depicted in Figure 1. Additional group-specific characteristics are described in detail elsewhere (Miles et al., 2018, 2019).

**TABLE 1.** SAMPLE CHARACTERISTICS. **Data presented as frequency counts or mean ± standard deviation.** AN = anorexia nervosa; BMI = body mass index

|  |  |
| --- | --- |
| n | 72 |
| Age, years | 28.3 ± 7.3 |
| BMI, kg/m<sup>2</sup> | 19.2 ± 3.0 |
| Education, years | 15.7 ± 1.8 |
| Race, white:non-white | 60:12 |
| Handedness, right:left | 66:6 |
| Lifetime AN diagnosis, yes:no | 48:24 |
| Subtype, restricting:binge-purge | 26:22 |
| Illness duration, years | 10.9 ± 8.1 |

**TABLE 2.** CLINICAL SCORES BY GROUP. **Data presented as mean ± standard deviation.** ^a^ vs. acAN, p(t)_HSD_ < 0.05; ^b^ vs. recAN, p(t)_HSD_ < 0.05; ^c^ vs. sibAN, p(t)_HSD_ < 0.05; ^d^ vs. HC, p(t)_HSD_ < 0.05; ^e^ includes drive for thinness, bulimia and body dissatisfaction subscales; acAN = women with acute anorexia; AN = anorexia nervosa; EDI-3 = Eating Disorder Inventory-3; HC = unrelated healthy controls; recAN = women with remitted anorexia; sibAN = unaffected sisters of acAN/recAN participants; STAI = State-Trait Anxiety Inventory; WCST-64:CV2 = WCST-64: Computer Version 2-Research Edition

| Measure | acAN | recAN | sibAN | HC |
| --- | --- | --- | --- | --- |
| Core AN symptoms (EDI-3) <sup>e</sup> | 57.9 ± 19.3 <sup>b,c,d</sup> | 36.6 ± 25.2 <sup>a,c,d</sup> | 16.1 ± 7.6 <sup>a,b</sup> | 14.1 ± 10.1 <sup>a,b</sup> |
| Trait anxiety (STAI) | 61.5 ± 13.4 <sup>b,c,d</sup> | 51.1 ± 13.1 <sup>a,c,d</sup> | 33.4 ± 6.1 <sup>a,b</sup> | 35.9 ± 11.8 <sup>a,b</sup> |
| Perseverative responses (WCST-64:CV2) | 5.9 ± 2.3 | 5.0 ± 2.2 | 4.9 ± 1.2 | 4.1 ± 0.6 |

**FIGURE 1.**
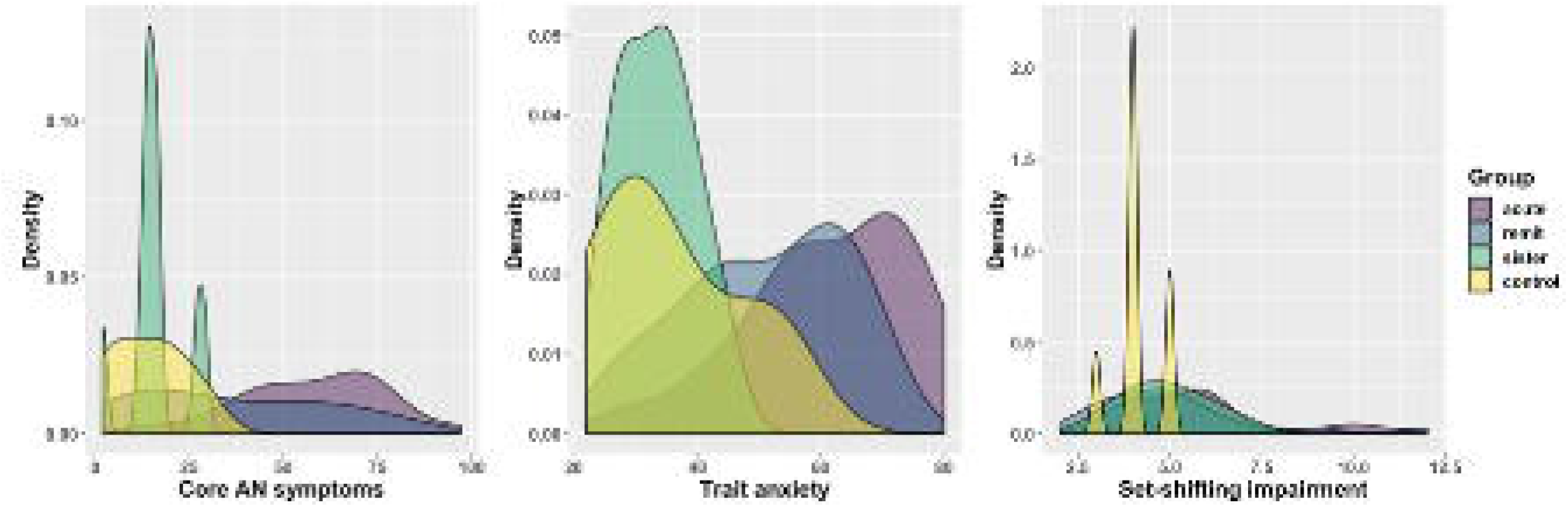
CLINICAL SCORE DISTRIBUTIONS. **Distributions of clinical scores in each participant subgroup.** AN = anorexia nervosa

We recruited seventy-two young adult women (28 ± 7 years), the majority of whom were non-Hispanic Caucasian, right-handed, and university-educated (Table 1). Two-thirds of our sample had a lifetime diagnosis of AN, and slightly more than half of those participants met criteria for restricting-subtype anorexia. Mean illness duration in women with acute or remitted AN was approximately eleven years.

As anticipated, clinical variables of interest varied widely across individuals and disease states (Fig. 1, Table 2). Consistent with previous studies in AN, core AN symptomatology and trait anxiety were significantly greater in women with a lifetime diagnosis of AN than in women without a lifetime diagnosis of AN, and they were greater during acute illness than in remission. Contrary to expectation, there was no main effect of group on set-shifting impairment.

### Regression results

Core AN symptomatology and trait anxiety showed distinct, spatially-distributed main effects on vertex-wise cortical thickness and LGI, while core AN symptomatology and set-shifting impairment showed partially overlapping effects on vertex-wise LGI. These results are summarized in Table 3 and depicted in Figure 2. We did not detect significant main effects of core AN symptomatology, trait anxiety, or set-shifting impairment on regional subcortical volume, vertex-wise cortical surface area or voxel-wise FA or MD.

**TABLE 3.** REGRESSION RESULTS. **Clusters in which clinical score is a significant predictor of cortical surface architecture, controlling for age, BMI and lifetime AN diagnosis.** ^a^ quantified using the drive for thinness, bulimia and body dissatisfaction subscales of the Eating Disorder Inventory-3; ^b^ quantified using the State-Trait Anxiety Inventory; ^c^ quantified using the perseverative responses score from the Wisconsin Card Sorting Test; AN = anorexia nervosa; Annot. = Desikan-Killiany atlas-based parcellation in which the peak vertex is located; CT = cortical thickness; DV = dependent variable; IV = independent variable; L = left hemisphere; LGI = local gyrification index; R = right hemisphere

| IV | DV | $t_{\max}$ | $p(t)_{\max}$ | Size (mm<sup>2</sup>) | Annot. |
| --- | --- | --- | --- | --- | --- |
| Core AN symptoms <sup>a</sup> | CT, cluster #1 | -4.521 | < 0.001 | 164 | lateral occipital, R |
| ''' | LGI, cluster #2 | -3.629 | < 0.001 | 4123 | insula, L |
| ''' | LGI, cluster #3 | -2.620 | < 0.001 | 3675 | superior frontal, R |
| ''' | LGI, cluster #4 | -2.997 | < 0.001 | 1959 | precentral, R |
| ''' | LGI, cluster #5 | -2.571 | < 0.001 | 895 | supramarginal, R |
| Trait anxiety <sup>b</sup> | CT, cluster #1 | -4.054 | < 0.001 | 130 | lingual, L |
| ''' | LGI, cluster #2 | -2.588 | 0.001 | 791 | superior parietal, L |
| ''' | LGI, cluster #3 | -2.284 | 0.001 | 858 | superior parietal, R |
| Set-shifting impairment <sup>c</sup> | LGI, cluster #1 | -3.687 | < 0.001 | 2415 | postcentral, R |

**FIGURE 2.**
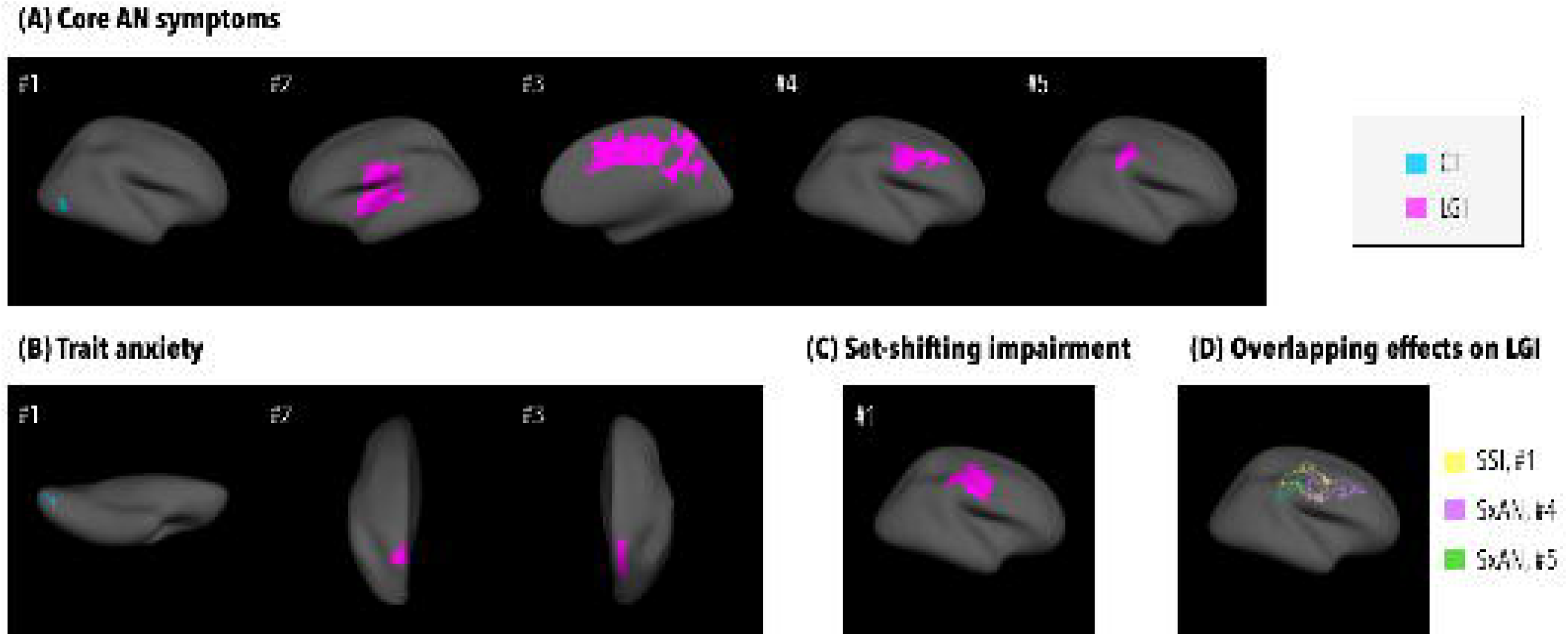
REGRESSION RESULTS. **Clusters in which clinical score is a significant predictor of cortical surface architecture, controlling for age, BMI and lifetime AN diagnosis.** (A)quantified using the drive for thinness, bulimia and body dissatisfaction subscales of the Eating Disorder Inventory-3; (B) quantified using the State-Trait Anxiety Inventory; (C) quantified using the perseverative responses score from the Wisconsin Card Sorting Test; AN = anorexia nervosa; CT = cortical thickness; LGI = local gyrification index; SSI = set-shifting impairment; SxAN = core AN symptoms

Greater core AN symptomatology was uniquely associated with lower cortical thickness in a right lateral occipital cluster (Fig. 2A, cluster #1) and with lower LGI in a left insular cluster and a right cingulate cluster (Fig 2A, clusters #2, #3). Likewise, greater trait anxiety was uniquely associated with lower cortical thickness in a left lingual cluster (Fig. 2B, cluster #1) and with lower LGI in bilateral superior parietal clusters (Fig. 2B, clusters #2, #3). Greater levels of core AN symptomatology and set-shifting impairment were associated with lower LGI in right sensorimotor and right inferior parietal clusters (Fig. 2B, clusters #4, #5; Fig. 2C, cluster #1). Although the latter effects overlapped spatially (Fig. 2D), they were statistically independent; the observed relationships between core AN symptomatology and LGI remained significant when set-shifting impairment was included as a nuisance variable (cluster #4, *t* = −2.697, p(*t*) = 0.009; cluster #5, *t* = −2.491, p(*t*) = 0.016), and the observed relationship between set-shifting impairment and LGI remained significant when core AN symptomatology was included as a nuisance variable (cluster #1, *t* = −2.745, p(*t*) = 0.008).

### Post hoc analyses

Results of post hoc Pearson correlation analyses are summarized in Table 4, and they are depicted in Figure 3. In each case, we observed directionally-consistent brain-behavior relationships in most, if not all, participant subgroups, and at least one within-group association was statistically significant. The latter was most often the case in women with remitted AN. However, at least one significant brain-behavior association was detected in each participant subgroup, and we may have been underpowered to detect additional within-group associations; in several cases, group-specific correlation coefficients failed to reach statistical significance although they were commensurate with or larger than those in the full sample.

**TABLE 4.** BRAIN-BEHAVIOR ASSOCIATIONS. **Sample-wide and group-specific Pearson correlations between clinical score and cluster-wise cortical surface architecture.** ^a^ quantified using the drive for thinness, bulimia and body dissatisfaction subscales of the Eating Disorder Inventory-3; ^b^ quantified using the State-Trait Anxiety Inventory; ^c^ quantified using the perseverative responses score from the Wisconsin Card Sorting Test; *p*(*r*) ≤ 0.05 (bolded), *p*(*r*) ≤ 0.10 (bolded and italicized); acAN = women with acute anorexia; AN = anorexia nervosa; CT = cortical thickness; DV = dependent variable; HC = unrelated healthy controls; IV = independent variable; LGI = local gyrification index; recAN = women with remitted anorexia; sibAN = unaffected sisters of acAN/recAN participants

| IV | DV | $r_{\text{sample}}$ | $r_{\text{acAN}}$ | $r_{\text{recAN}}$ | $r_{\text{sibAN}}$ | $r_{\text{HC}}$ |
| --- | --- | --- | --- | --- | --- | --- |
| Core AN symptoms <sup>a</sup> | CT, cluster #1 | **-0.362** | **-0.693** | **-0.457** | -0.459 | **-0.835** |
| "" | LGI, cluster #2 | **-0.439** | -0.333 | **-0.488** | -0.422 | -0.189 |
| "" | LGI, cluster #3 | **-0.367** | **-0.420** | **-0.475** | -0.491 | 0.370 |
| "" | LGI, cluster #4 | **-0.397** | **-0.381** | **-0.486** | -0.482 | 0.300 |
| "" | LGI, cluster #5 | **-0.374** | **-0.443** | -0.312 | **-0.678** | 0.139 |
| Trait anxiety <sup>b</sup> | CT, cluster #1 | **-0.288** | 0.005 | **-0.656** | **-0.563** | -0.583 |
| "" | LGI, cluster #2 | **-0.331** | -0.358 | **-0.562** | -0.245 | 0.104 |
| "" | LGI, cluster #3 | **-0.259** | -0.037 | **-0.610** | -0.488 | -0.141 |
| Set-shifting impairment <sup>c</sup> | LGI, cluster #1 | **-0.414** | **-0.414** | **-0.465** | -0.107 | -0.613 |

**FIGURE 3.**
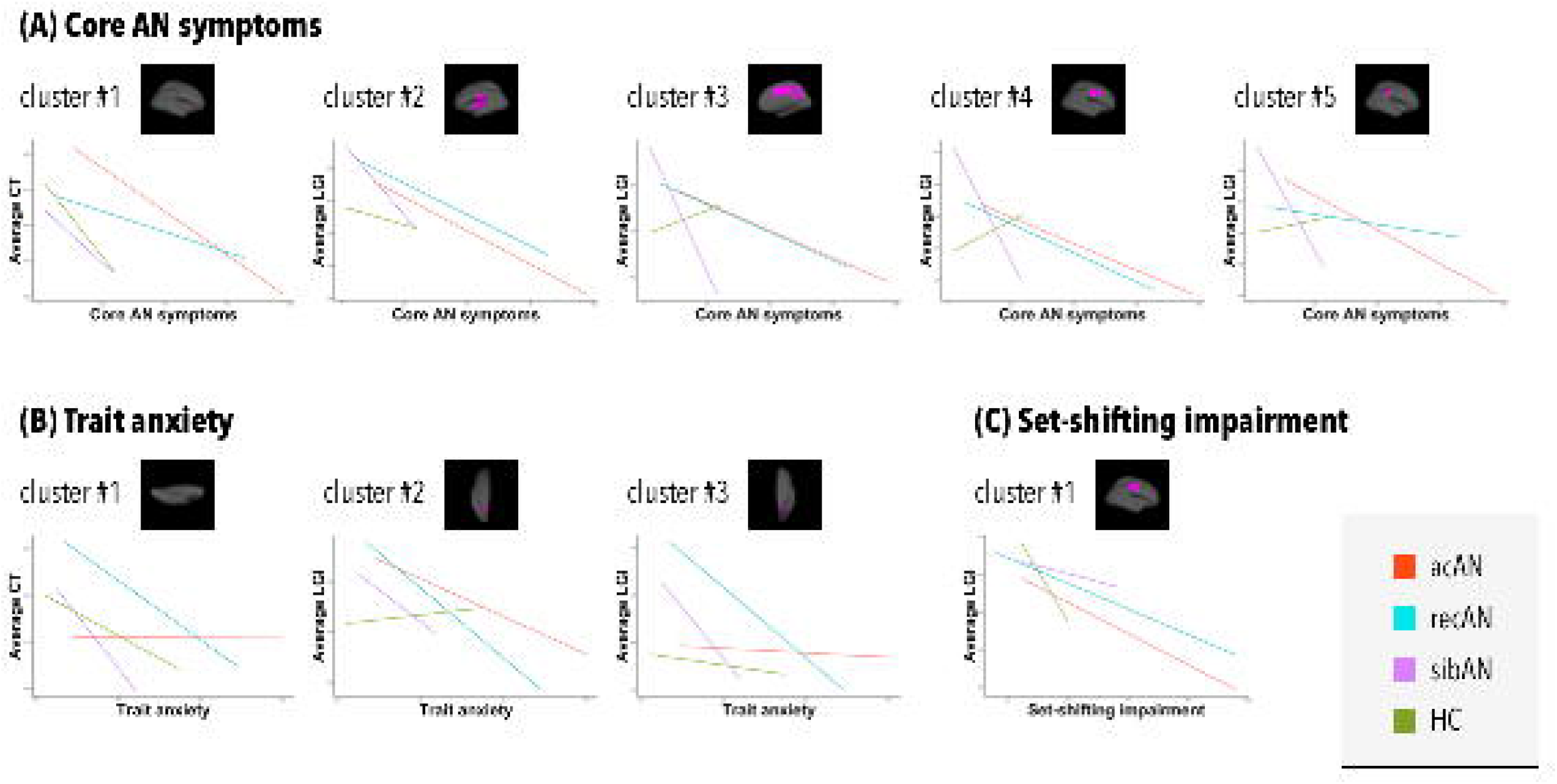
BRAIN-BEHAVIOR ASSOCIATIONS. **Group-specific correlations between clinical score and cluster-wise cortical surface architecture.** acAN = women with acute anorexia; AN = anorexia nervosa; CT = cortical thickness; HC = unrelated healthy controls; LGI = local gyrification index; recAN = women with remitted anorexia; sibAN = unaffected sisters of acAN/recAN participants

Cross validation analyses provided additional evidence of generalizability as model fit remained relatively good when random subsets of participants were excluded (RMSE = 0.113 – 0.285) and when each participant subgroup was excluded (RMSE = 0.116 – 0.306).

## DISCUSSION

In this study, we sought to identify novel grey and white matter signatures of core features of AN and shared temperamental and cognitive-behavioral risk factors for AN by dimensionally mapping relationships between distinct neuroanatomical phenotypes and clinical variables of interest. In order to reduce the likelihood that observed relationships were driven by malnutrition effects, we tested brain-behavior associations in a mixed sample, including women with acute and remitted AN, their unaffected sisters, and unrelated healthy controls, and we controlled for current BMI and lifetime AN diagnosis.

As anticipated, we detected significant negative associations between measures of core AN symptomatology and trait anxiety and site-specific cortical thickness, and we detected significant negative associations between each clinical measure of interest and site-specific local gyrification. Contrary to expectation, however, we detected no significant associations between clinical variables of interest and regional subcortical volume, site-specific cortical surface area, or site-specific white matter microstructure.

### Distinct associations between core AN symptomatology and site-specific cortical surface architecture

We detected a significant negative association between core AN symptomatology and cortical thickness, a highly heritable component of cortical volume, thought to represent number of neurons within a cortical column, that peaks in middle childhood and decreases linearly across adult life (Hogstrom, Westlye, Walhovd & Fjell, 2013; Panizzon et al., 2009; Rakic, 1988; Raznahan et al., 2011), in a cluster covering part of the right lateral occipital cortex. This finding partially mirrors the negative association between drive for thinness and right extrastriate cortical thickness reported by King et al. (2015). However, whereas this group reported a state-dependent link, we replicated the aforementioned association in each subgroup, excluding unaffected sisters. As such, our finding is consistent with previous reports of site-specific cortical thinning and symptom exacerbation in acute illness, and it provides novel evidence of a link between site-specific cortical thickness and disease risk.

The lateral occipital cortex, part of the extrastriate visual association cortex, plays a key role in object recognition (Grill-Spector, Kourtzi & Kanwisher, 2001), and it forms part of the ventral and lateral visual networks. Given that age-related decline is slower in the occipital lobe than in frontal, temporal, or parietal lobes (Hogstrom et al., 2013), low cortical thickness in this region could reflect accelerated cortical thinning in adolescence and early adulthood, perhaps driven by a potentially-heritable variation in neurodevelopmental programming. Greater cortical thinning could, in turn, increase vulnerability to AN by disrupting communication between and/or within networks subserving perception of and/or attention to body shape and size. Altered resting-state functional connectivity, consistent with this hypothesis, has been reported in AN; Phillipou et al. (2016) reported reduced resting-state functional connectivity between visual association and sensorimotor areas in women with acute AN, and Favaro et al. (2012) and Scaife, Godier, Filippini, Harmer & Park (2017) reported reduced resting-state functional connectivity within ventral and lateral visual networks, respectively, in women with acute and remitted AN.

We also detected significant negative associations between core AN symptomatology and LGI, a moderately heritable index of cortical folding (Rogers et al., 2010), the patterning of which is thought to optimize connectivity between adjacent regions (Klyachko & Stevens, 2003), that peaks in early life and remains relatively stable across adult life (Armstrong, Schleicher, Omran, Curtis & Zilles, 1995; Cao et al., 2017), in clusters covering parts of the left insular cortex and right cingulate cortex. This finding is partially consistent with evidence of site-specific hypogyrification in AN (Bernardoni et al., 2018; Leppanen et al., 2019). Unlike group effects from case-control studies, however, our sample-wide findings replicated most strongly in women with remitted AN, suggesting a dimensional link between site-specific local gyrification and disease risk that is trait-based, not state-dependent.

The insular cortex and the cingulate cortex are highly interconnected regions that subserve diverse cognitive, emotional, sensorimotor, and homeostatic functions, including attention, interoception, and emotional experience (Taylor, Seminowicz & Davis, 2009; Uddin, Nomi, Hébert-Seropian, Ghaziri & Boucher, 2017). As part of broader functional networks, including the default mode and cingulo-opercular networks (Sadaghiani & D’Esposito, 2014), these regions have been implicated in subjective self-representation (Taylor et al., 2009), a process that involves integrating interoceptive and emotional salience information, and maintenance of cognitive control (Dosenbach et al., 2007). Low LGI in these regions could reflect reduced cortical folding in early life, possibly resulting from potentially-heritable variations in neurodevelopmental programming. Less cortical folding could, in turn, increase vulnerability to AN by disrupting communication between and/or within networks subserving self-concept formation and appropriate response selection (Medford & Critchley, 2010), particularly in the context of energy homeostasis. Again, altered resting-state functional connectivity, consistent with this hypothesis, has been reported in AN; Ehrlich et al. (2015) reported reduced resting-state functional connectivity within a thalamo-insular subnetwork in acute AN, and Gaudio, Olivo, Zobel & Schiöth (2018) reported reduced resting-state functional connectivity within a cerebellar-insular-parietal-cingular subnetwork in acute AN. Of note, the latter study compared adolescent girls at the earliest stages of AN to age-matched healthy controls, reducing the likelihood that observed effects were products of chronic malnutrition.

Beyond disrupting communication within or between networks, site-specific hypogyrification could contribute to more nuanced patterns of atypical connectivity. For example, it could facilitate concurrent increases and decreases in resting-state functional connectivity between the anterior cingulate cortex and parts of the default mode network (Lee et al., 2014) and executive control network (Gaudio et al., 2015), respectively, in acute AN. Disproportionate communication between the anterior cingulate cortex and default mode regions could give rise to body and food-related rumination while insufficient communication between the anterior cingulate cortex and executive control regions could drive cognitive and behavioral rigidity that reinforces food avoidance.

### Overlapping associations between core AN symptomatology and set-shifting impairment and site-specific local gyrification

Cognitive and behavioral rigidity, while not unique to eating disorders, has been proposed as an endophenotype of AN (Holliday, Tchanturia, Landau, Collier & Treasure, 2005). Therefore, it is not surprising that we detected overlapping, yet independent, associations between core AN symptomatology and set-shifting impairment and LGI in a right frontoparietal region. This finding is partially consistent with evidence of state-dependent, site-specific hypogyrification in AN (Miles et al., 2018), although replication in women with remitted AN and/or unaffected sisters suggests trait-relevance.

As part of the frontoparietal control network, the aforementioned region has been implicated in task-set initiation and performance feedback (Dosenbach et al., 2007), processes that are crucial for flexible coordination of goal-driven behavior. Given evidence of significant genetic correlations between AN and OCD (Anttila et al., 2016), greater overall genetic risk for AN could contribute to more developmental hypogyrification in adjacent and overlapping frontoparietal regions. Hypogyrification across more of the frontoparietal cortex could, in turn, increase both vulnerability to AN and potential severity of AN by driving multi-dimensional deficits in cognitive and behavioral flexibility that manifest as body and food-related rumination and persistent food avoidance and hinder adaptive information processing necessary for treatment engagement. Reduced resting-state functional connectivity, consistent with this hypothesis, has been reported in remitted AN (Boehm et al., 2016), and it has also been observed in OCD (Guersel, Avram, Sorg, Brandl & Koch, 2018). Compounding effects of variations in cortical surface architecture associated with core AN symptomatology and set-shifting impairment could contribute to worse outcomes observed in AN patients with more pronounced cognitive-behavioral deficits (Steinhausen, 2002).

### Distinct associations between trait anxiety and site-specific cortical surface architecture

Cortical thinning in the left lingual cortex and hypogyrification in the bilateral superior parietal cortex, both of which were associated with greater trait anxiety, could likewise contribute to AN severity. Cortical thinning has been reported in major depressive disorder (Na et al., 2016), high rates of which have been reported in AN (Halmi et al., 1991), and hypogyrification in the left superior parietal cortex has been associated with trait anxiety in healthy adults (Miskovich et al., 2016). Moreover, in a longitudinal study by Favaro, Tenconi, Degortes, Manara & Santonastaso (2015), lower gyrification in the left superior parietal cortex distinguished acute AN patients and controls at baseline, and it distinguished patients with good outcome at follow up from those with poor outcome at follow up.

The lingual cortex has been implicated in visual memory (Bogousslavsky, Miklossy, Deruaz, Assal & Regli, 1997), and the superior parietal cortex has been implicated in visual perception, spatial cognition, reasoning, working memory, and attention (Wang et al., 2016). Given the aforementioned evidence of disruption in AN within resting-state networks subserving similar functions (i.e., visual, default mode, and frontoparietal control networks), the combination of AN- and anxiety-associated variations in cortical surface architecture could give rise to complex clinical presentations and contribute to poorer outcomes observed in AN patients with higher levels of general psychopathology (Herpertz◻Dahlmann, Wewetzer, Schulz & Remschmidt, 1996).

### Limitations and future directions

We tested relationships between measures of core AN symptomatology, trait anxiety, and set-shifting impairment and distinct neuroanatomical phenotypes in a sample enriched for AN vulnerability. With fewer than one-hundred participants, we may have been underpowered to detect subtle brain-behavior relationships. Likewise, by testing associations across subjects, rather than within groups, we may have failed to detect brain-behavior relationships that are state-dependent. Ideally, replication would occur in the context of a longitudinal multimodal MRI study, including an equal number of unaffected sisters and healthy controls and an additional group of patients with anxiety-spectrum disorders and/or OCD, that is sufficiently powered to model complex relationships among structural and functional brain phenotypes (i.e., cortical surface architecture and resting-state functional connectivity) and clinical variables of interest and can examine the impact of subtype, BMI, and illness duration in AN patients.

Despite limitations in sample size and composition, this study complements those designed to test group differences in brain structure, and it provides key additional evidence of brain-based risk for AN. Specifically, it provides evidence of (1) distinct associations between core AN symptomatology and cortical thickness in a right lateral occipital region and local gyrification in left insular and right cingulate regions; (2) distinct associations between trait anxiety and cortical thickness in a left lingual region and local gyrification in bilateral superior parietal regions; and (3) overlapping associations between core AN symptomatology and set-shifting impairment and local gyrification in a right frontoparietal region. Perhaps most importantly, it does so while controlling for lifetime AN and current BMI, the most prominent feature of acute illness, and its sample-wide findings are largely replicated within-group, reducing the likelihood that results are best attributed to malnutrition.

In sum, our findings suggest that site-specific variations in cortical surface architecture, some of which have been observed in previous case-control studies in AN, contribute to core AN symptomatology and shared temperamental and cognitive-behavioral risk factors for AN. Moreover, our study suggests that the aforementioned variations in brain structure form part of a vulnerability profile that predates disorder development and could influence prognosis.

## ACKNOWLEDGEMENTS AND DISCLOSURES

The authors would like to sincerely thank staff at the CAMH Research Imaging Centre and members of the Kimel Family Imaging-Genetics Laboratory, particularly Dr. Erin Dickie, for their assistance with MRI acquisition and processing.

## Notes

This work was supported by the CAMH AFP Innovation Fund [2014, 2016].

Dr. Miles, Dr. Nikolova, and Dr. Voineskos report no conflicts of interest. Dr. Kaplan has received honoraria and lecture fees from Shire Pharmaceuticals.

